# Valvular Prostaglandins are Elevated in Severe Human Aortic Valve Stenosis

**DOI:** 10.1101/2023.08.14.553317

**Authors:** Lucien G.J. Cayer, Arun Surendran, Tobias Karakach, Harold M. Aukema, Amir Ravandi

## Abstract

**Introduction:** Aortic valve stenosis (AVS) is the most common valvular disease in the developed world. AVS involves the progressive fibro-calcific remodeling of the aortic valve (AV), which impairs function and can ultimately lead to heart failure. Due to gaps in our understanding of the underlying mechanisms of AVS, there are no pharmacological treatments nor dietary interventions known to slow AVS progression. Recent studies have begun to suggest oxylipins, a class of bioactive lipid, may be dysregulated in the valves of AVS-patients.

**Methodology:** We utilized HPLC-MS/MS to conduct a targeted oxylipin analysis on human AV tissue and plasma from a cohort of 110 patients undergoing AV surgery.

**Results:** We identified 36 oxylipins in human AV tissue with all showing significant increase in patients with severe AVS. A multivariate model including patient characteristics and valvular oxylipins identified arachidonic acid-cyclooxygenase (COX) pathway derived prostanoids to be the most associated with AVS severity. Plasma oxylipin levels were measured in a subset of aortic surgery patients and compared to a control group of healthy participants, showing distinct oxylipin profiles between control and disease.

**Conclusion:** Our comprehensive analysis of oxylipins in the human AV to date and identified the inflammatory and osteogenic regulating prostanoids to be positively correlated with AVS severity. This elucidation of prostanoid dysregulation warrants further research into COX inhibition to mitigate AVS.

## Introduction

Aortic Valve Stenosis (AVS) is a debilitating cardiovascular condition that affects millions of individuals worldwide. In Western nations, it is the third most common cardiovascular disease, the most common valvular heart disease, and a major cause of heart failure and cardiovascular mortality ^1–3^. The progression of AVS is characterized by fibro-calcific remodelling of the aortic valve (AV) leaflets which result in a narrowing of the valve opening and diminished function ^4^. Risk factors of AVS include age, elevated low-density lipoprotein, smoking history, hypertension, and diabetes ^5,6^. There is currently no effective pharmacological intervention for AVS, and surgical valve replacement or transcatheter AV implantation are the current best treatment methods, but they may not be feasible for patients with significant comorbidities ^7^. There is a growing need to better understand the underlying mechanisms of AVS to develop medical treatments and dietary recommendations, both to reduce the incidence and progression, while minimizing the need for invasive therapies.

AVS was previously thought to be a passive disease associated with aging and wear-and-tear of the valve but has since been shown to be an active inflammatory process. The AV leaflet is comprised primarily of valvular interstitial cells (VICs) that with AVS develop an osteoblast-like phenotype ^8^. This process is initiated at points of mechanical stress on the AV leaflet, which incur infiltration of immune cells and oxidized lipoproteins which occur in early lesions ^4,9,10^. Oxidized lipoproteins are cytotoxic and propagate further inflammation. The inflammatory cytokines released, stimulate fibrosis via fibroblast differentiation into myofibroblasts. Calcification occurs with the presence of macrophages in the initial stages of inflammation, but is accelerated due to continued release of cytokines promoting differentiation of myofibroblasts to osteoblast-like cells ^8^. This in turn stimulates the production of pro-calcific proteins (osteopontin, osteocalcin, osteonectin and bone morphogenetic protein-2 & –4). This process results in mineral deposition in the extracellular matrix resulting in bone-like tissue within the leaflet that is supported by angiogenesis ^4,11^. It has recently been hypothesized that statin-therapy would reduce the risk of AVS, similarly to atherosclerosis, by reducing lipoproteins. Unfortunately, large-scale prospective randomized control clinical trials have failed to show beneficial results from multiple intensive statin-therapies ^12–15^. There is currently no effective pharmacological treatment for AVS, thus further elucidation of the mechanism of AVS is required.

Oxylipins are a class of lipid mediators which are produced by enzymatic oxidation of polyunsaturated fatty acids (PUFA). It has now been shown that enzymes involved in oxylipin synthesis, namely phospholipase A2, cyclooxygenase (COX)-1, COX-2, and 5-lipoxygenase (LOX), are elevated in calcified areas of AV leaflet tissue ^16–19^. Further, a recent study has identified two specialized pro-resolving mediators, the eicosapentaenoic acid (EPA) derived resolvin (Rv)E1 and docosahexaenoic acid (DHA) derived RvD3 to be lower, while leukotriene (Lt)B4 was elevated, in the calcified regions of the AVS leaflet ^20^. However, like the endothelial cells or whole heart, the lipidomic profile of AV leaflets is complex and we anticipate that there is a larger panel of bioactive oxylipins that more accurately characterize calcification of AV leaflets. ^21,22^.

The progression of AVS is, in part, driven by inflammatory mechanisms that resemble those of maladapted tissue repair. In general, the arachidonic acid (AA) derived oxylipins, such as the prostanoids, mediate proinflammatory cellular processes, while the n-3 PUFA derived oxylipins, which include the specialized pro-resolving mediators, mediate anti-inflammatory and tissue repair processes ^23,24^. The dysregulation of inflammation-regulating oxylipins should consequently be expected in AVS. Further, oxylipins mediate cellular functions other than inflammation. Of particular relevance to AVS are prostaglandin (PG)E2 and PGF2α due to their involvement in the process of osteogenesis, which plays a critical role in the development of the disease ^25–28^. Thus, there is a need to compare the oxylipins in calcified and non-calcified human aortic valves to further our understanding of the underlying pathophysiology of AVS. Here we quantify for the first time the whole profile of oxylipins in the human AV and plasma of patients receiving aortic surgery and identify the alterations in oxylipin profiles with AVS severity.

## Methods

### Human aortic valve tissue

The full details of the participant recruitment and tissue collection have been previously reported ^29^. Research ethics boards at the University of Manitoba and St. Boniface Hospital approved collection of these tissues for biomolecular study. Briefly, male and female aortic surgery patients (n=110) at St. Boniface Hospital (Winnipeg, Canada) were recruited and provided written informed consent. AV mean transvalvular gradient (MPG) and calcification severity were based on transthoracic echocardiography measurements ^30^. Pre-surgical plasma samples were also collected from a subset of patients (n=17). The AV leaflets were surgically removed and immediately placed in an ice-cold ethylenediaminetetraacetic acid/phosphate buffered saline solution, and then flash frozen for storage at –80°C.

Plasma was obtained from an additional group of patients with moderate to severe AVS based on echocardiographic assessment (n=55) and a group of healthy control patients with no history of AVS (n=44). A complete table of characteristics for the control patients can be found in Supplemental Table 1.

### Oxylipin extraction and quantification

For this study, free oxylipins were extracted and quantified from AV leaflets as described in ^31,32^, with modifications. Briefly, whole leaflets were weighed and homogenized in Tyrode’s salt solution (9.6 g/L, pH 7.6) at a ratio of 7 mg to 40 µL, using an Omni Bead Ruptor 24 (Omni International, Inc., GA, USA). Antioxidant cocktail (0.2 g/L butylated hydroxytoluene, 0.2 g/L ethylenediaminetetraacetic acid, 2 g/L triphenylphosphine, and 2 g/L indomethacin in methanol/ethanol/water (2:1:1, v/v/v)) was added (1:20, v/v) to the homogenates, followed by the addition (4:3, v/v) of methanol/formic acid (100:1, v/v). A mixture of deuterated oxylipin internal standards (Cayman Chemical, MI) was added to the AV homogenates. Samples were diluted in acidified water (1:5, v/v) to adjust to pH 3 and applied to 33 μm polymeric reverse phase extraction columns (Phenomenex, CA, USA) pre-conditioned with methanol and pH 3 water. The columns were rinsed with pH 3 water followed by hexane. Oxylipins were eluted from the column with methanol, collected, evaporated, and resuspended in MS grade water: acetonitrile: acetic acid (70:30:0.02; v/v/v) for high performance liquid chromatography tandem mass spectrometry (Shimadzu Nexera XR coupled to a QTRAP 6500 triple quadruple; Sciex, ON) analysis, by a procedure modified from ^33^ and described in ^31,34^. Oxylipins were quantified using the stable isotope dilution method described in ^35^, such that the oxylipin concentration was calculated from the ratio of the peak area, at an established retention time and mass-to-charge ratio, to that of a known amount of deuterated internal standard. Oxylipin concentrations were reported in relation to total valvular protein determined by extraction with RIPA lysis buffer followed by bicinchoninic acid protein assay.

### Data analyses

Clinical parameters and oxylipins were compared by ANOVA, using AVS severity groups. Patients were grouped into AVS severity groups using MPG (mild <20 mmHg, moderate 20-40 mmHg, and severe ≥40 mmHg, based on the 2014 AHA/ACC guidelines ^36^), or by calcification score (mild: 1-2, moderate: 3, and severe 4-5). Data was transformed using Tukey’s ladder of powers to achieve normality followed by ANOVA and Tukey’s HSD test. When normality was not achieved, the Kruskal-Wallis test was used with Dunn’s post hoc test. The Bonferroni correction for multiple comparisons was then applied. Initially sex was included as a factor in the linear model but was removed due to an absence of significant sex-effects for any oxylipin. Linear correlations between oxylipins and clinical parameters were determined using Pearson’s correlation test. Principal component analysis (PCA) was performed on per sample unit sum normalized clinical data and log_10_ oxylipins. A stepwise linear regression model was also performed to identify which oxylipins and clinical parameters independently were predictive of MPG. All statistical analyses were performed using R (version 4.2.2) implemented through RStudio (version 2023.03.1+446) for graphical visualization.

## Results

### Patient characteristics

The baseline characteristics of the patients grouped by MPG severity from which valve samples were are displayed in Table 1. Age, sex, height, body surface area, blood pressure, blood lipids, blood glucose, creatinine, hypertension history, medications, and bicuspid aortic valve did not differ significantly between severity groups. Only weight and body mass index differed between groups, with a significant increase with AVS severity. Smoking history was highest among the moderate group. Calcification and echocardiograph parameters predictably worsened with MPG severity. Further details have been previously reported in ^29^.

**Table 1:**
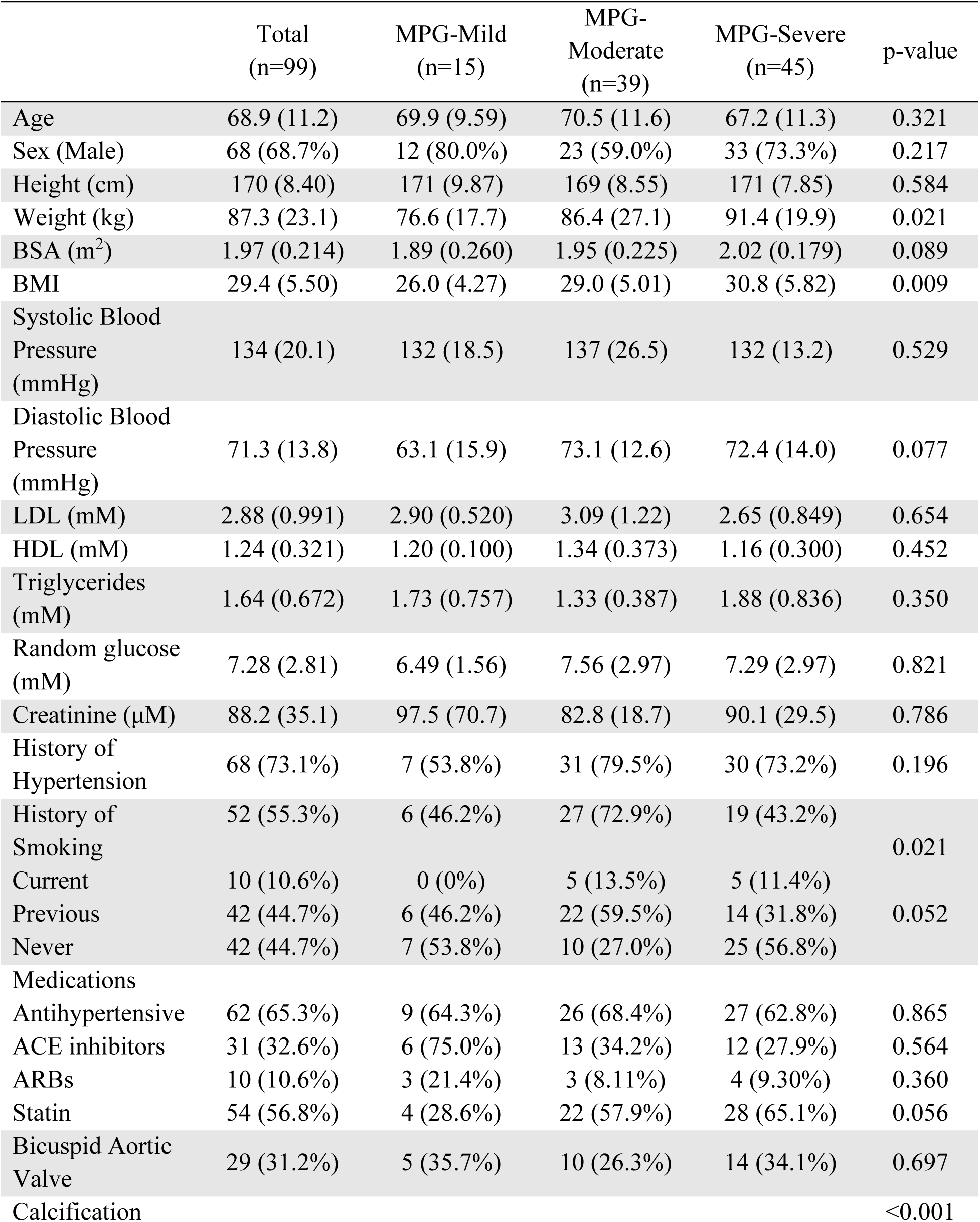
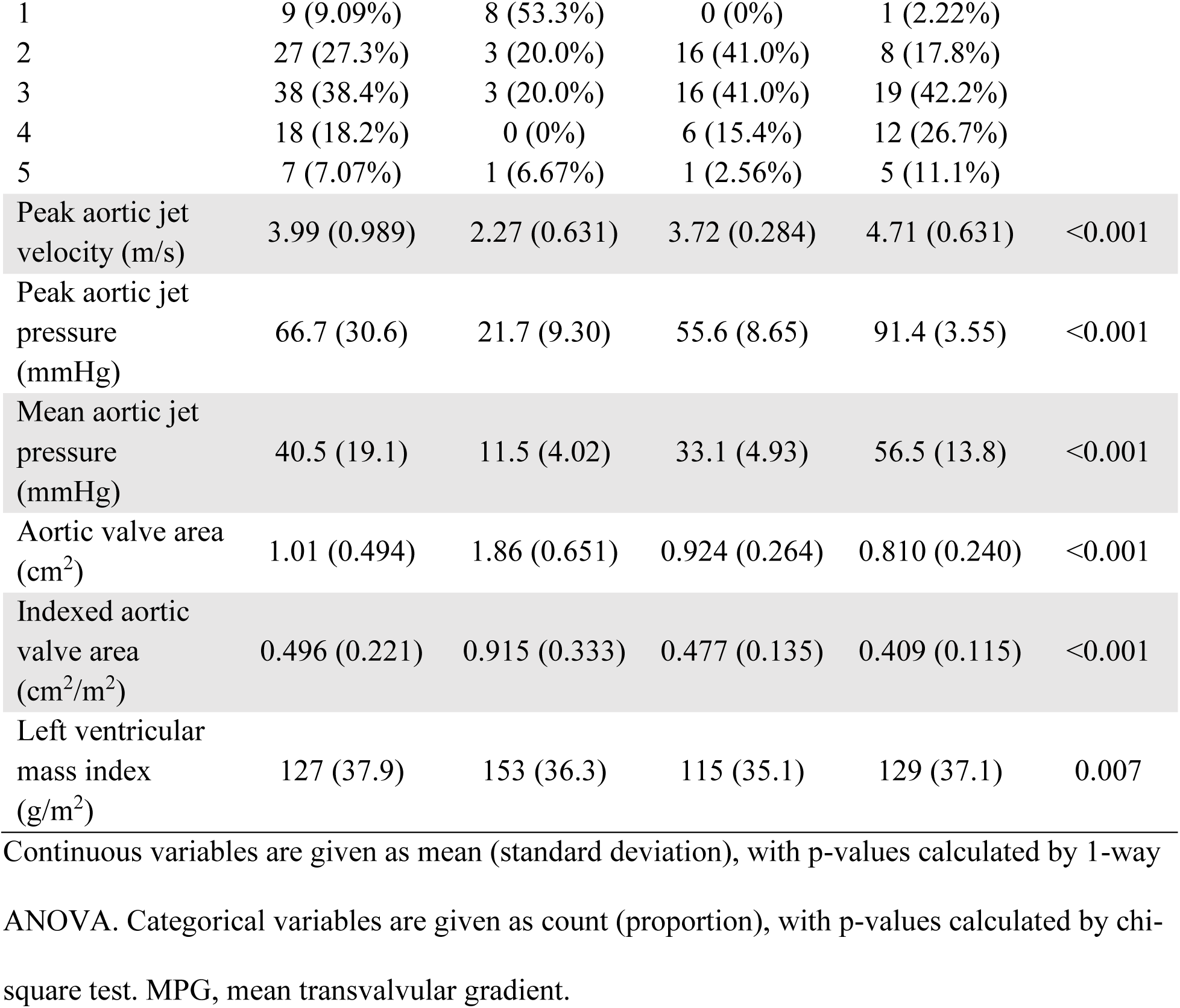
Patient characteristics for aortic valve oxylipin analysis.

Table 2 displays the baseline patient characteristics for the plasma oxylipins analysis. There were notable differences between the healthy control and AVS groups. The control group was younger, with lower rates of current and past smoking, and had lesser use of ACE inhibitors or ARBs and statins.

**Table 2:**
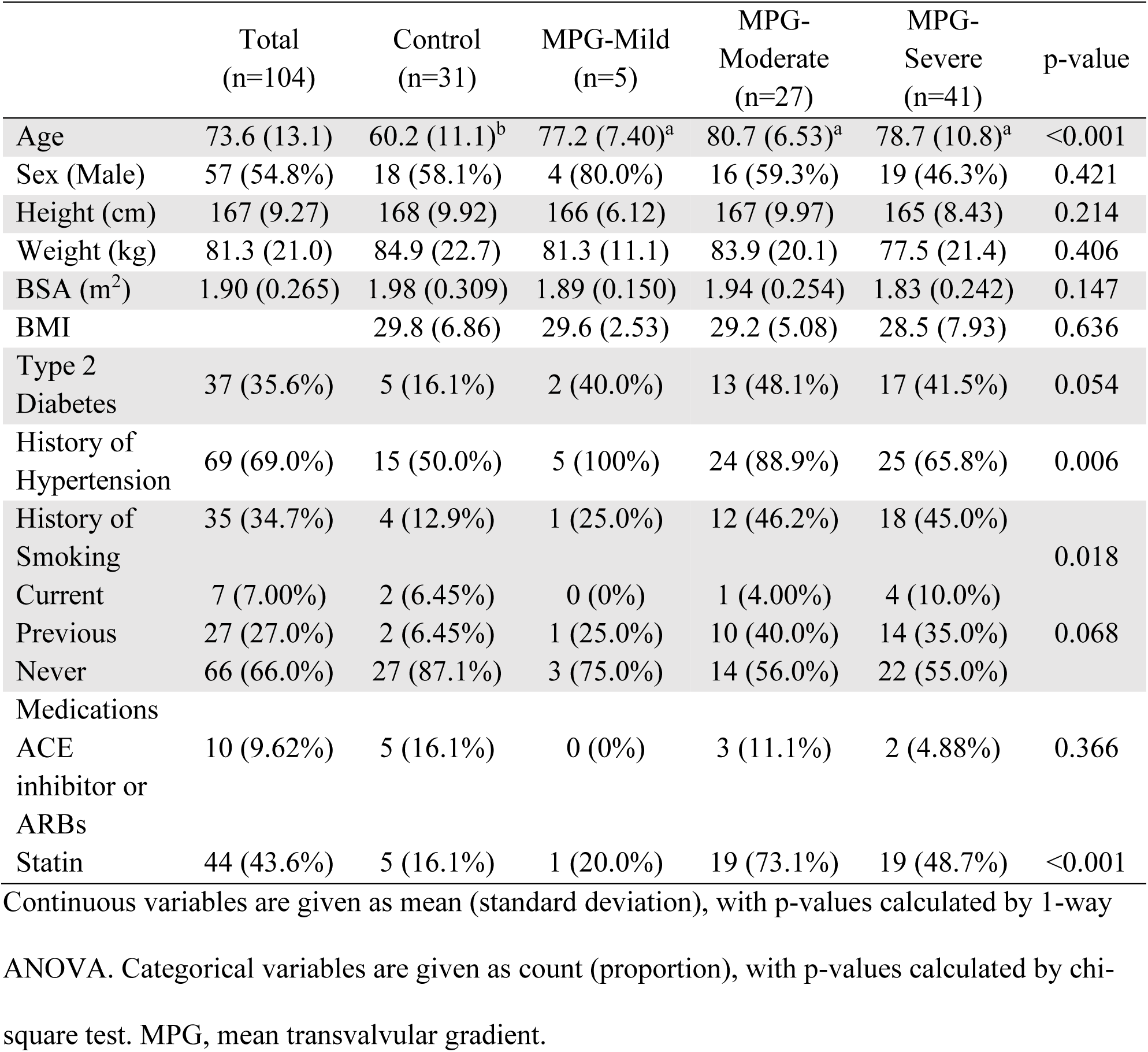
Shared patient characteristics for plasma analysis.

### Valvular oxylipins

We quantified 36 unique oxylipins in valvular tissue. The complete list of the oxylipins quantified and comparisons by MPG severity can be found in Supplemental Table 2. Total oxylipin levels were found to increase with disease severity, and this was reflected in the total oxylipin mass by MPG (Fig. 1A). When patients were grouped by MPG severity, total oxylipins was elevated when comparing MPG mild to moderate and mild to severe (Fig. 1B). Similarly, the total oxylipins increased when patients were grouped by calcification score (Fig. 1C). Each oxylipin quantified increased with AVS severity (Fig. 2A). From the volcano plots, we see 33 and 30 out of 36 oxylipins had a log_2_-fold change greater than 1 and a p-value less than the Bonferroni adjustment for multiple comparisons (*α=0.0014*) when comparing MPG mild to severe and calcification mild (1,2) to. severe (4,5), respectively (Fig. 2B, C).

**Figure 1.**
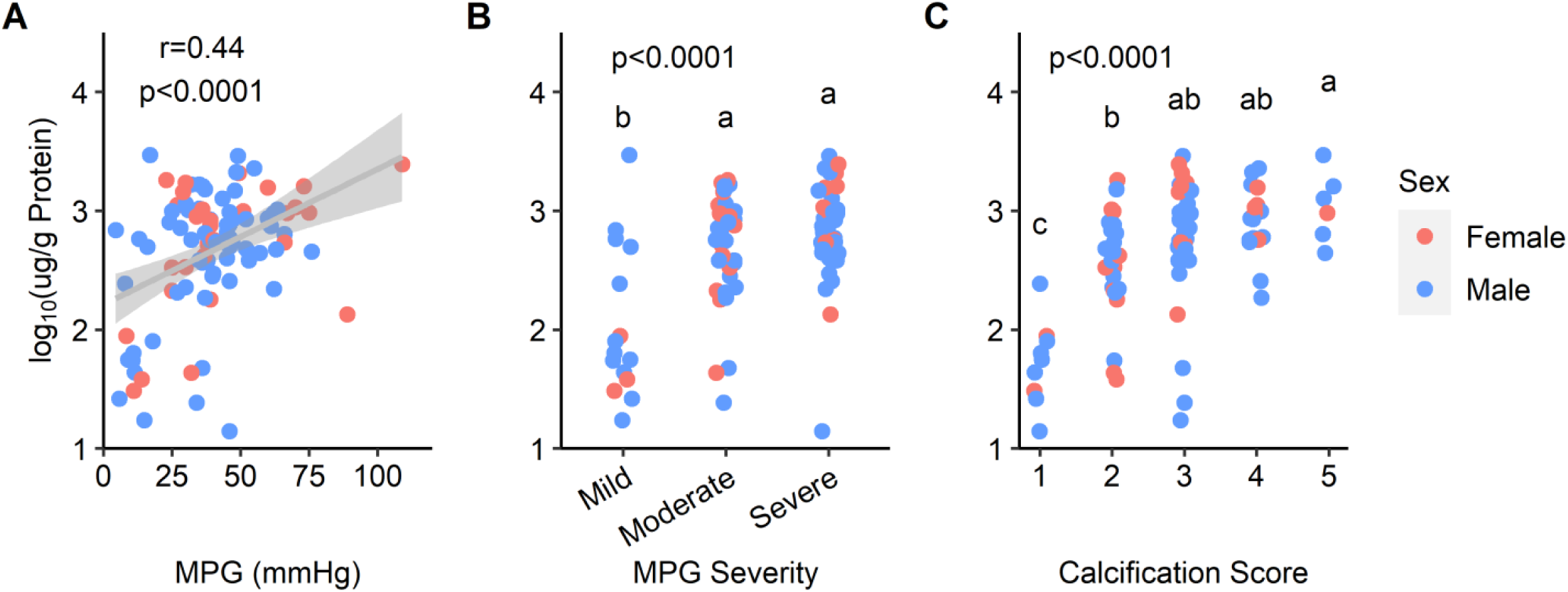
A) Total aortic valve oxylipins by mean pressure gradient. Trendline has a 95% confidence interval. **B) Total aortic valve oxylipins by MPG severity. C) Total aortic valve oxylipins by calcification score.**

**Figure 2.**
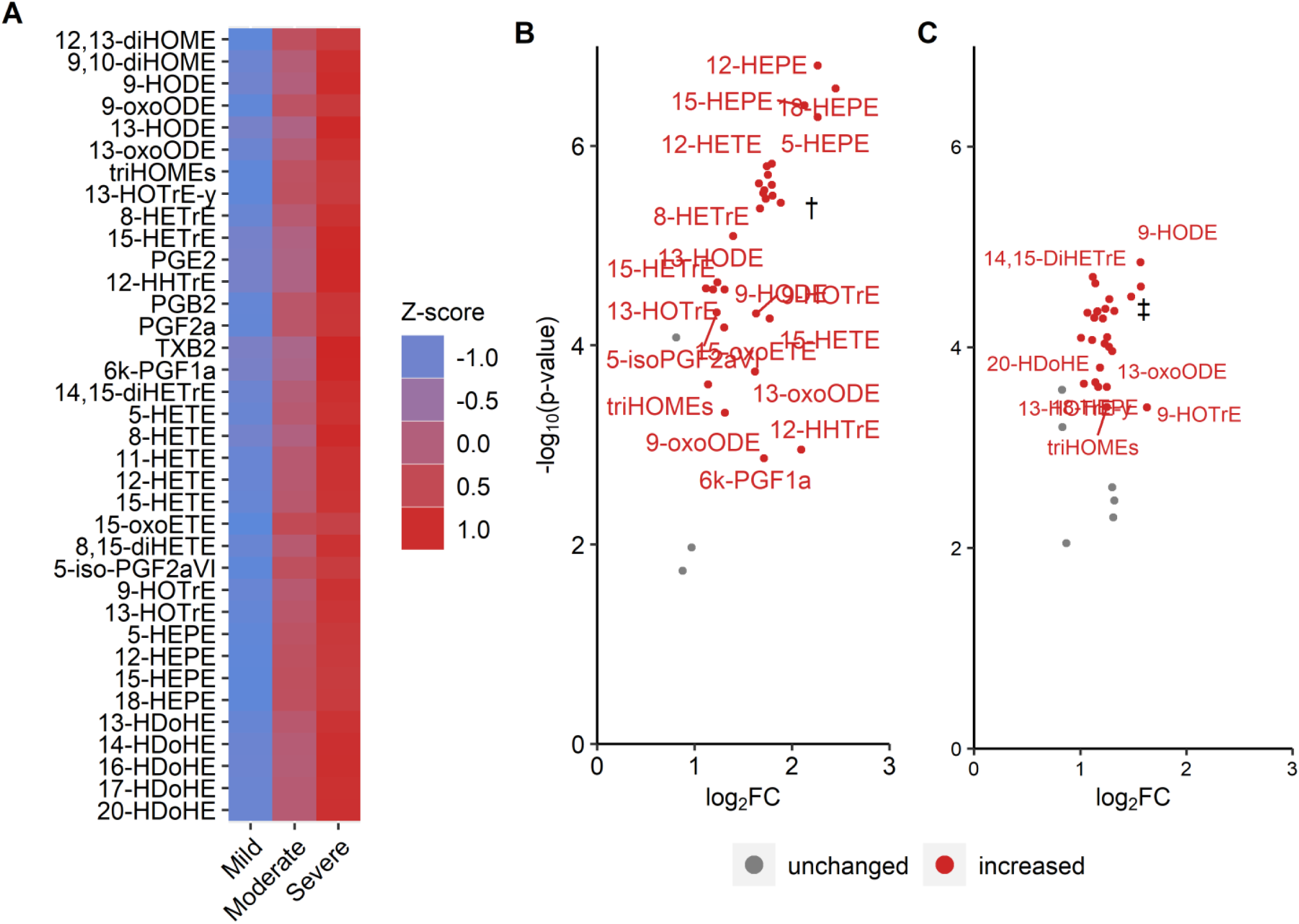
A) Heatmap of aortic valve oxylipins by mean transvalvular gradient severity. Values are normalized for each oxylipin with mean centering and root-mean-square scaling. **B) Volcano plot of valvular oxylipins, MPG: mild vs. severe.** Oxylipins were considered up-regulated if log2FC > 1 (severe relative to mild AVS), and p-value <0.0014 (Bonferroni adjusted α). **C) Volcano plot of valvular oxylipins, calcification score: mild (1,2) vs. severe (4,5).** Oxylipins were considered up-regulated if log2FC > 1 (severe relative to mild AVS), and p-value <0.0014 (Bonferroni adjusted). Abbreviations: diHETE, dihydroxy-eicosatetraenoic acid; diHETrE, dihydroxy-eicosatrienoic acid; diHOME, dihydroxy-octadecaenoic acid; HDoHE, hydroxy-docosahexaenoic acid; HEPE, hydroxy-eicosapentaenoic acid; HETE, hydroxy-eicosatetraenoic acid; HETrE, hydroxy-eicosatrienoic acid; HHTrE, hydroxyheptadecatrienoic acid; HODE, hydroxy-octadecadienoic acid; HOTrE, hydroxy-octadecatrienoic acid; oxoETE, oxo-eicosatetraenoic acid; oxoODE, oxo-octadecadienoic acid; PG, prostaglandin; triHOME, trihydroxy-octadecenoic acid (9,10,13-triHOME & 9,12,13-triHOME peaks could not be separated); TX, thromboxane.

When oxylipins levels were categorized based on the fatty acid precursor from which they are derived, each of the fatty acid precursors was elevated with AVS severity whether dividing patients by MPG or calcification severity (Fig. 3). Specifically, the moderate and severe groups were significantly elevated from mild, but not distinct from each other. When the fatty acid totals were considered as proportions of the total oxylipins, we see that the relative amount of linoleic acid oxylipins decreased with between MPG mild and severe, while AA, α-linolenic acid, dihomo-γ-linolenic acid, and EPA oxylipins increased. For calcification severity, only the increased proportion of ALA reached significance. When oxylipins were totalled by the major enzymatic pathways from which they were derived, each pathway was increased with AVS severity (MPG and calcification), such that moderate and severe groups were significantly elevated from mild, but not distinct from each other (Fig. 4). For enzyme pathways, the proportions of the total oxylipins did not change with AVS severity.

**Figure 3.**
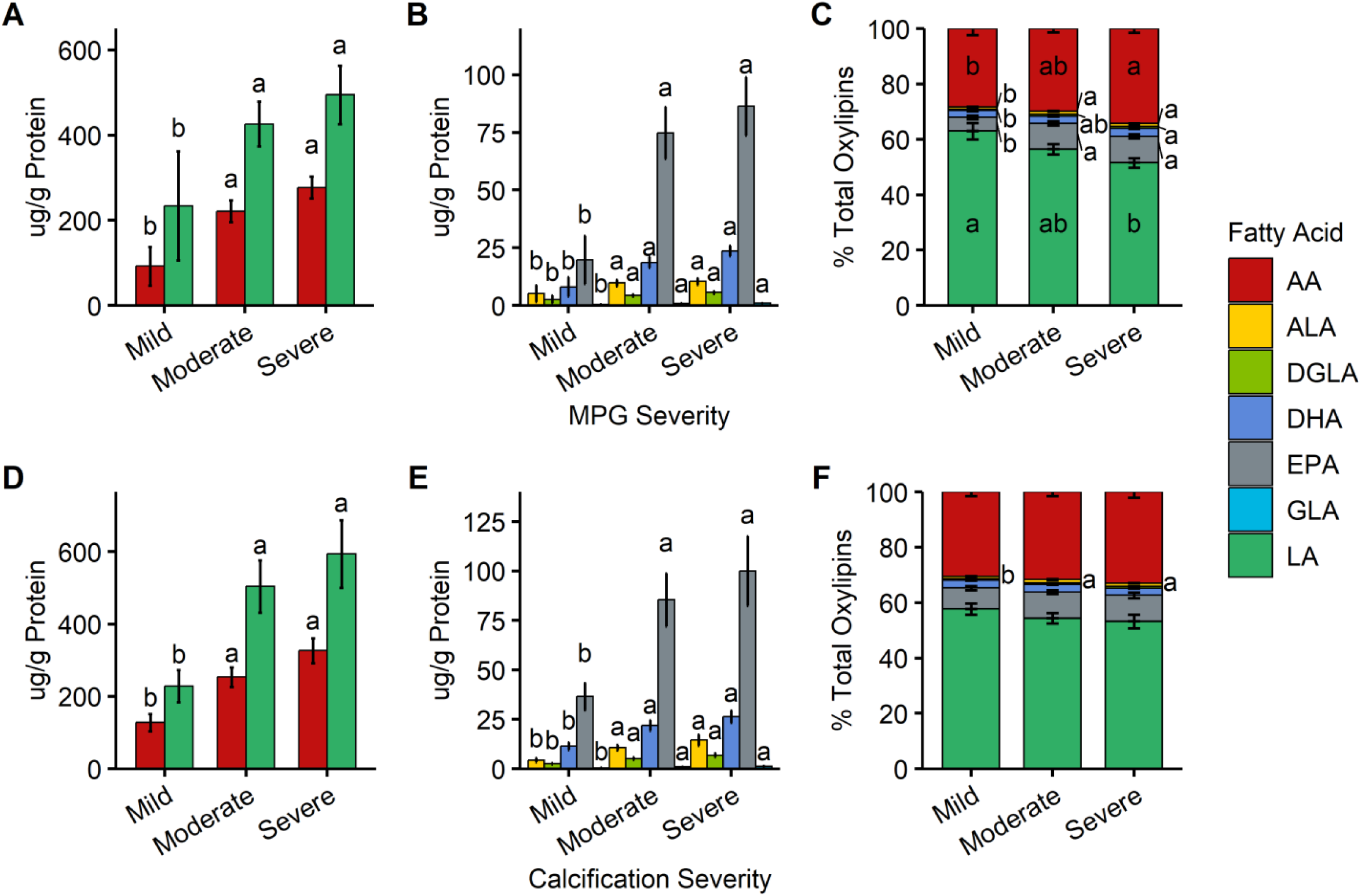
Aortic valve oxylipins totalled by fatty acid precursor: A-C) MPG severity. D-F) Calcification severity. Comparisons are between severity groups and within each fatty acid precursor. Abbreviations: AA, arachidonic acids; ALA, α-linolenic acid; DGLA, dihomo-γ-linolenic acid; DHA docosahexaenoic acid; EPA, eicosapentaenoic acid; GLA, γ-linolenic acid; LA, linoleic acid; MPG, mean transvalvular pressure gradient.

**Figure 4.**
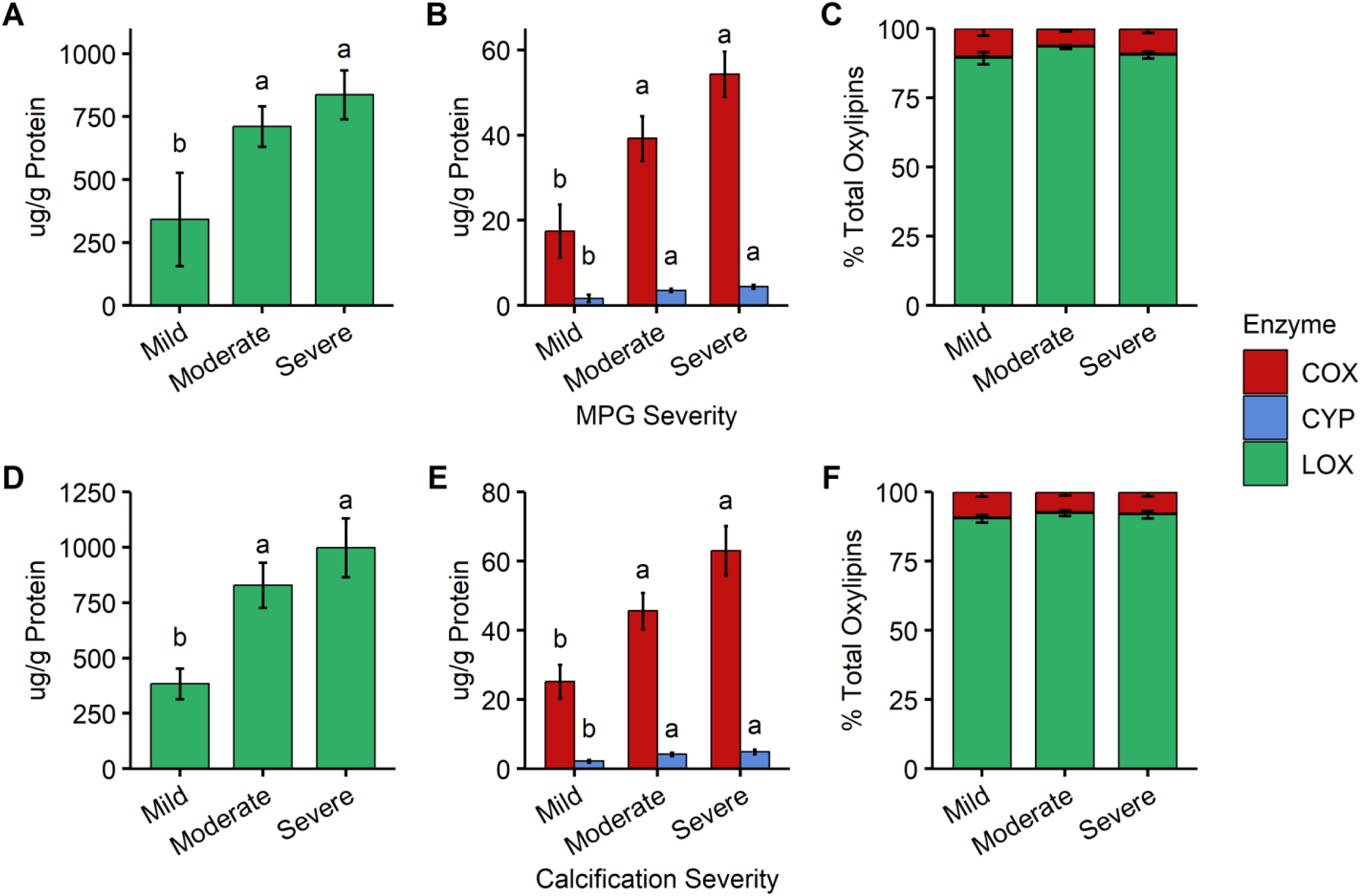
Aortic valve oxylipins totalled by enzymatic pathway: A-C) MPG severity. D-F) Calcification severity. Comparisons are between severity groups and within each enzymatic pathway. Abbreviations: COX, cyclooxygenase; CYP, cytochrome P450; LOX, lipoxygenase; MPG, mean transvalvular pressure gradient.

### Valvular oxylipins and clinical parameters

Figure 5 depicts the Pearson correlations between each of the valvular oxylipins and clinical parameters. There are distinct groupings of highly correlated variables, such as the AVS severity parameters. Also, the majority of oxylipins are highly correlated with each other; notable exceptions were PGE2, 12-hydroxy-heptadecatrienoic acid, PGF2α, thromboxane (TX)B2, and 6-keto-PGF1α, which are best correlated with each other and are all derived from the AA-COX pathway.

**Figure 5.**
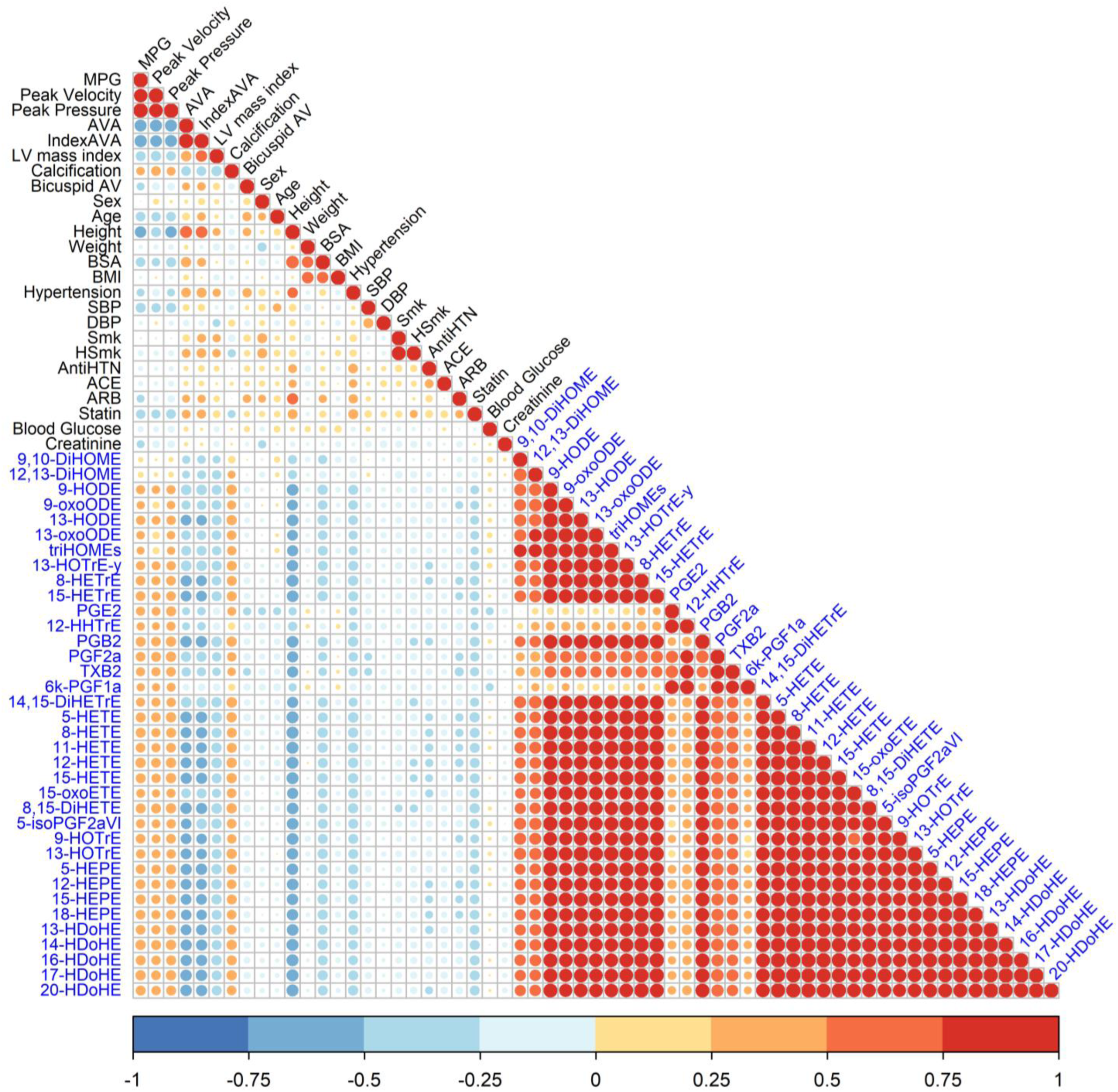
Pearson correlations of valvular oxylipins and clinical parameters. Variables in black text are clinical parameters and variables in blue text are oxylipins. Abbreviations: AntiHTN, antihypertensive; AV, aortic valve; AVA, aortic valve area; DBP, diastolic blood pressure; DiHETE, dihydroxy-eicosatetraenoic acid; DiHETrE, dihydroxy-eicosatrienoic acid; DiHOME, dihydroxy-octadecaenoic acid; HDoHE, hydroxy-docosahexaenoic acid; HEPE, hydroxy-eicosapentaenoic acid; HETE, hydroxy-eicosatetraenoic acid; HETrE, hydroxy-eicosatrienoic acid; HHTrE, hydroxyheptadecatrienoic acid; HODE, hydroxy-octadecadienoic acid; HOTrE, hydroxy-octadecatrienoic acid; HSmk, smoking history; IndexAVA, indexed aortic valve area; LV, left ventricle; MPG, mean transvalvular pressure gradient; oxoETE, oxo-eicosatetraenoic acid; oxoODE, oxo-octadecadienoic acid; PG, prostaglandin; SBP, systolic blood pressure; Smk, current smoking; triHOMEs, trihydroxy-octadecenoic acid; TX, thromboxane.

Next, a multivariate model, that included all oxylipins and clinical parameters was constructed. From PCA, figure 6A depicts the scores plot, in which each patient is projected on to components 1 and 2. The AVS severity status has been overlayed onto each patient and AVS severity can be seen to be positively associated with each component. Figures 6B and C display the top absolute loadings values for variables on components 1 and 2, respectively. These show that component 1 primarily accounts for the variance in the model attributed to the majority of oxylipins, while the variables contributing to component 2 include AVS severity markers and AA COX-derived prostanoids. Specifically, MPG, peak aortic jet velocity, peak aortic jet pressure, PGE2, PGF2α, 6k-PGF1α, 12-hydroxy-heptadecatrienoic acid, and TXB2 were positively associated, and AV area, indexed AV area, and left ventricle mass index – which are inversely related to increased AVS severity – were negatively associated. Thus, this multivariate model shows that prostanoids in the aortic valve are highly correlated with AVS severity. These findings are validated by a multiple linear regression model with MPG as the dependant variable that was optimized by stepwise selection and resulted in the inclusion of the clinical parameters (age, weight, systolic blood pressure, and creatinine) and PGF2α as independent variables (Table 3).

**Figure 6.**
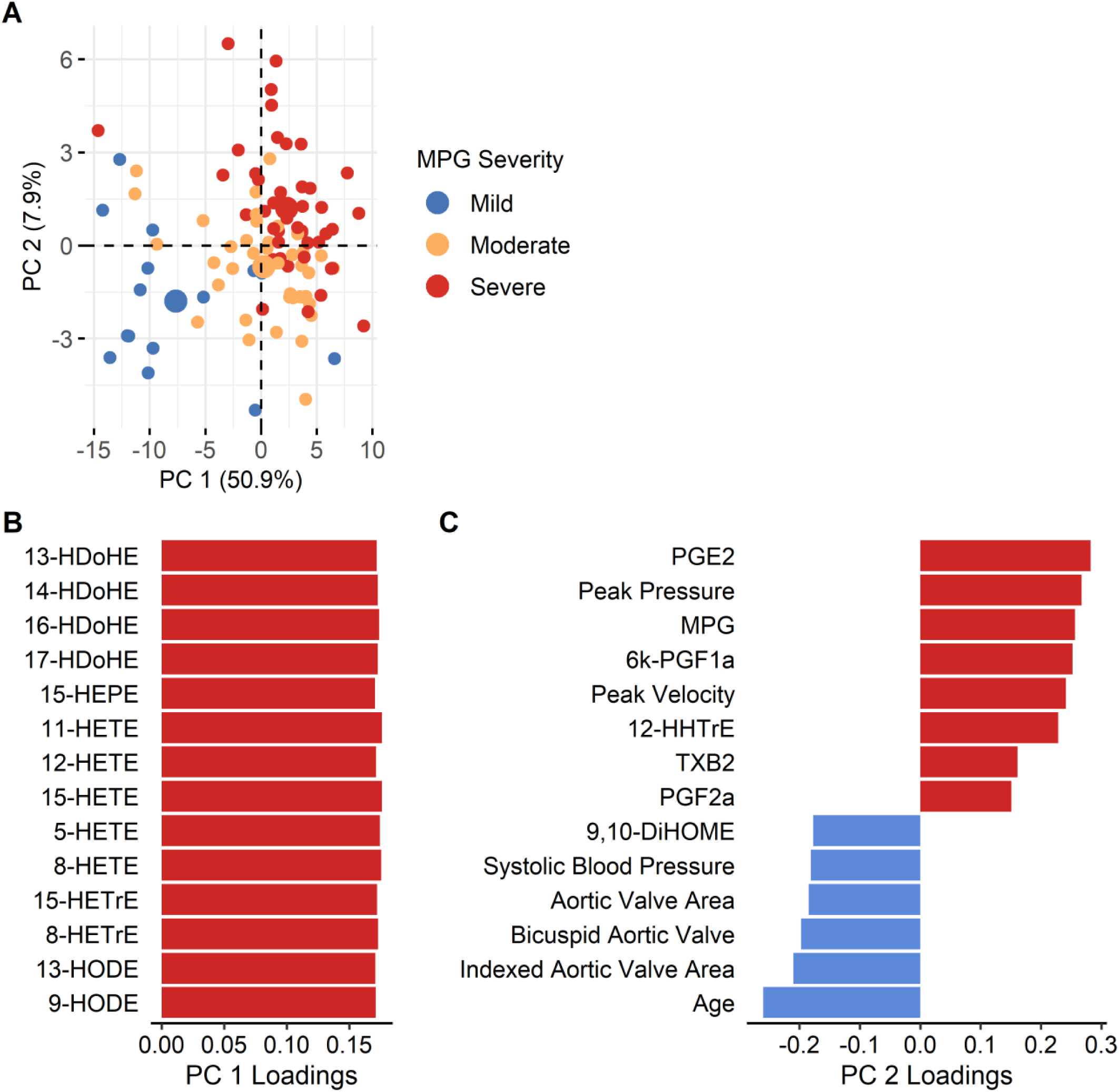
Patient characteristics and valvular oxylipins modeled by principal component analysis. **A) Scores plot of individuals on principal components 1 and 2.** Colour overlayed by MPG severity group. **B) Loadings values of the top variables by absolute value for component 1. C) Loadings values of the top variables on PC 2.** Abbreviations: DiHOME, dihydroxy-octadecaenoic acid; HDoHE, hydroxy-docosahexaenoic acid; HEPE, hydroxy-eicosapentaenoic acid; HETE, hydroxy-eicosatetraenoic acid; HETrE, hydroxy-eicosatrienoic acid; HHTrE, hydroxyheptadecatrienoic acid; HODE, hydroxy-octadecadienoic acid; MPG, mean transvalvular pressure gradient; PC, principal component; PG, prostaglandin; TX, thromboxane.

**Table 3.** Effect of clinical parameters and oxylipins on aortic valve mean transvalvular gradient.

|  | Estimate | Std. Error | t-value | Pr(>\|t\|) |
| --- | --- | --- | --- | --- |
| (Intercept) | 0.153 | 0.0157 | 9.74 | <0.0001 |
| Age | -0.400 | 0.108 | -3.71 | 0.0003 |
| Weight | -0.181 | 0.0663 | -2.73 | 0.0076 |
| SBP | -0.319 | 0.0692 | -4.61 | <0.0001 |
| Creatinine | -0.197 | 0.0438 | -4.50 | <0.0001 |
| PGF2α | 6.95 | 2.27 | 3.06 | 0.0029 |

Abbreviations: PG, prostaglandin; SBP, systolic blood pressure.

### Plasma oxylipins

Plasma oxylipins from patients with severe AVS were compared to control healthy participants without AVS (Supplemental Table 4). Total oxylipins were found to be higher in the severe AVS patients compared to controls (Fig. 7A). Unlike the valvular oxylipins, dysregulation of plasma oxylipins occurred in both directions, with both up and down-regulation with disease severity (Fig. 7B). The oxylipin totals by fatty acid precursors were compared: AA, DHA, γ-linolenic acid oxylipins were decreased and α-linolenic acid, dihomo-γ-linolenic acid, DPA, EPA, and linoleic acid oxylipins increased in AVS patients (Fig. 7C). When compared as proportions of the total oxylipin mass all oxylipin classes except EPA were significantly different between control and severe AVS patients (Fig. 7D).

**Figure 7.**
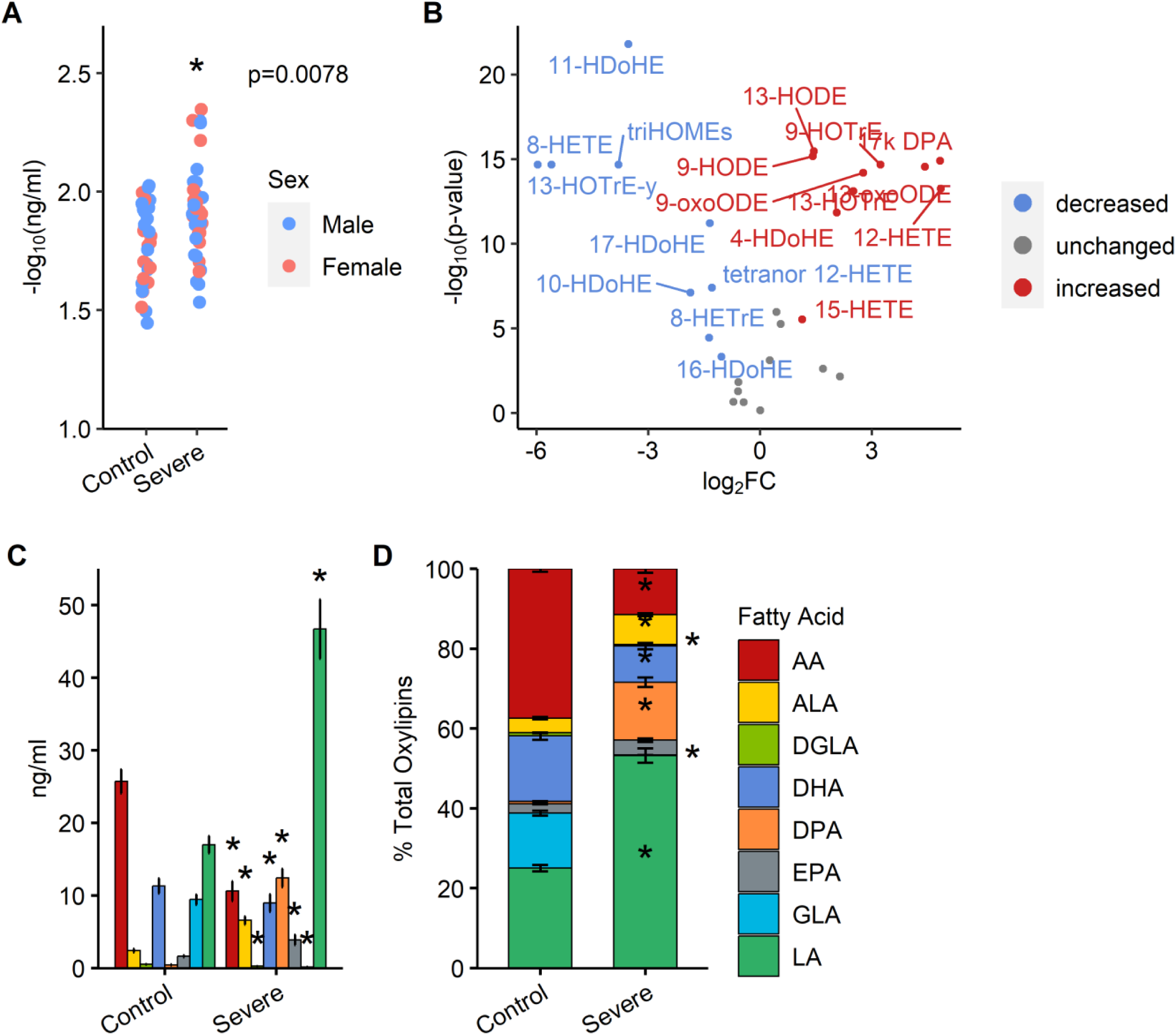
A) Total plasma oxylipins for control and MPG-severe patients. B) Volcano plot of plasma oxylipins, control vs. severe. Oxylipins were considered up-regulated if log2FC > 1 and p-value <0.0014 (Bonferroni adjusted α). **C-D) Plasma oxylipins by fatty acid precursor.** Comparisons between control and severe for each precursor PUFA. Abbreviations: AA, arachidonic acid; ALA, α-linolenic acid; DGLA, dihomo-γ-linolenic acid; DHA, docosahexaenoic acid; DPA, docosapentaenoic acid; EPA, eicosapentaenoic acid; GLA, γ-linolenic acid; LA, linoleic acid.

A PCA model that included the plasma oxylipins and clinical parameters of the control participants and patients from each AVS severity group shows clear separation between control and AVS patients along PC1 (Fig. 8A). The top absolute loadings values for variables on component 1 of this model are displayed in figure 8B. When we removed the controls from the model the is no separation based on AVS severity (Fig. 8C).

**Figure 8.**
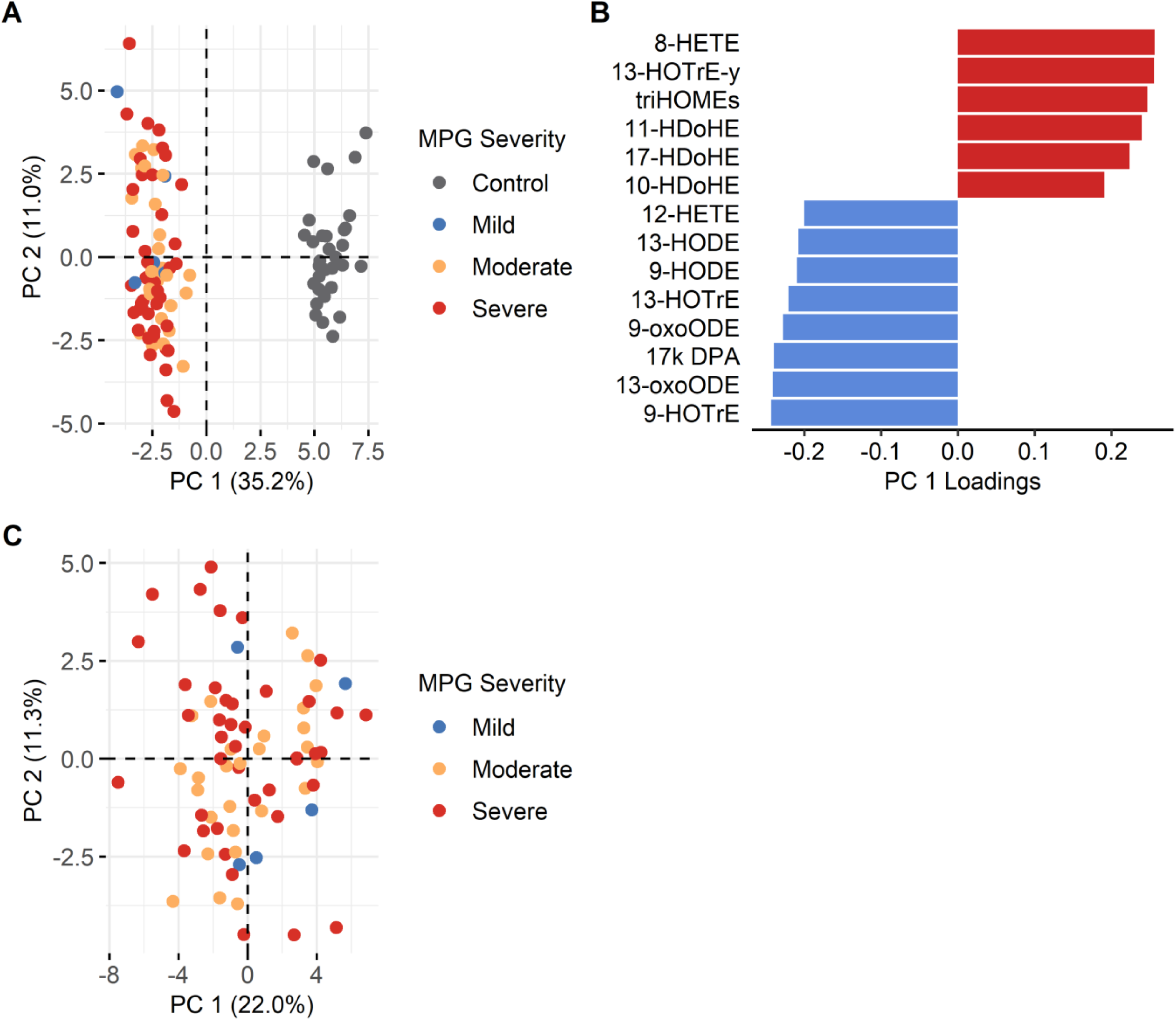
A-C) Patient characteristics and plasma oxylipins modeled by principal component analysis. A) Scores plot, all plasma sample patients on components 1 and 2 with colour overlayed by MPG severity group; B) Loadings values of the top variables on principal component 1. C) Scores plot from model excluding control patients. Abbreviations: DPA, docosapentaenoic acid; HDoHE, hydroxy-docosahexaenoic acid; HETE, hydroxy-eicosatetraenoic acid; HODE, hydroxy-octadecadienoic acid; HOTrE, hydroxy-octadecatrienoic acid; oxoODE, oxo-octadecadienoic acid; triHOME, trihydroxy-octadecenoic acid (9,10,13-triHOME & 9,12,13-triHOME peaks could not be separated).

## Discussion

This study provides the first comprehensive targeted analysis of the oxylipin profile in calcified human AV tissue. Our results indicate a broad increase in oxylipins with increasing AVS severity, suggesting a significant disruption of physiological homeostasis occurs. The dysregulation of oxylipins may contribute to the progression of AVS, as these lipid mediators play crucial roles in inflammation and osteogenesis.

The alterations in the valvular oxylipin profile, particularly the increase in the proportion of valvular AA oxylipins (Fig. 2C) and the identification of the AA-COX prostanoids as variables of importance by the multivariate modelling (Fig. 5C, Table 3), are informative to the mechanistic understanding of the chronic inflammation characteristic of AVS. The prostanoids are inflammatory mediators which play key roles in the initiation of inflammatory response ^37,38^. There is also evidence that prostanoids are involved in chronic inflammatory states by promoting persistent cellular infiltration by immune cells ^39^. Thus, our study supports the current mechanism and builds upon it by quantifying the elevation of specific prostanoids with AVS severity.

In addition to the contribution of inflammatory processes to AVS development, prostanoids are likely to contribute to AV calcification. Of the five prostanoids highlighted in this analysis, two have direct functionality connections to osteogenesis. Both PGE2 and PGF2α are mediators of calcium resorption and bone remodelling. PGE2 modulates the differentiation of myofibroblasts, osteoblasts, and osteoclasts ^40–43^; while, PGF2α regulates proliferation and differentiation of osteoblasts and their mineralization of Ca^2+^ and inorganic phosphate ^27,28^. 6k-PGF1α was also identified in this analysis, and it is a stable hydrolysis product of the short-lived prostacyclin (PGI2). There is emerging evidence that PGI2 may have osteogenic properties. A PGI2 synthase knockout mouse model had increased bone formation, resorption, and bone mass ^44^. Also, the stable PGI2 analog Iloprost improved bone regeneration in a mouse osteotomy trial ^45^. Similarly, TXB2 is a stable hydrolysis product of TXA2, which may induce the formation of osteoclasts ^46^. The effect of these PGs specifically in VICs has not yet been demonstrated, but these molecules are likely candidates for mediating the connection between COX up-regulation and the development of the osteoblast-like phenotype and mineral deposition that is characteristic of AVS progression. Thus, these data support and enhance the current understanding of the mechanisms of AVS development with the chronic inflammatory environment promoting calcification mediated through valvular prostanoids.

Currently, there is conflicting data on the role of COX in AVS. Firstly, COX-1 and COX-2 mRNA and protein expression have been found to be elevated in *ex-vivo* calcified AV tissues and *in-vitro* calcification models ^17–19^; while, one study conflictingly found decreased COX-2 expression *ex-vivo* ^47^. Our study’s findings of increased COX derived oxylipins supports the former. If we accept that COX expression increases with AVS, whether increased COX promotes valvular calcification or is a product of it is not known. On the other hand, it should be noted that COX expression and activity level are not always positively correlated ^48^.

Studies of COX inhibition with AVS models have been conducted but provide ambiguous data. In a *Klotho*-deficient mouse model for AVS and in cultured porcine aortic VICs in osteogenic conditions, the use of celecoxib (selective COX-2 inhibitor) was shown to slow calcification ^18^. Also, this mouse model demonstrated that increased valvular COX-2 expression preceded calcification. Contradictorily, in *ex-vivo* human and *in-vitro* porcine VICs, calcification induced with treatment of transforming growth factor β1 or osteogenic media was enhanced with celecoxib ^47,49,50^. However, Vaidya et al. (2020) found that celecoxib reduced calcification *in-* vitro when glucocorticoids present in osteogenic media were removed, concluding that the increased calcification observed was due to an interaction between glucocorticoids and not COX-2 inhibition ^50^.

Two retrospective analyses of observational studies have looked at AVS and COX inhibitors with one finding an association between selective COX-2 inhibitors and increased AVS risk and the other finding no association ^49,51^. Both studies also found non-selective COX inhibitors did not influence AVS risk ^49,51^. It is also worth noting that selective COX-2 inhibitors have been associated with an increased risk of cardiovascular adverse events ^52^.

There have been fewer studies that focussed on the COX-1 isozyme. Recently, Sakaue et al. (2020) found the elevation of COX-1 in calcified valvular tissue to be greater than that of COX-2 and identified COX-1 activity as being essential to the differentiation of VICs to the osteoblast-like phenotype *in-vitro* ^19^. Yet, in an observational study no association between AVS risk and aspirin (selective COX-1 inhibitor) was found, and a potential increased risk was shown in one cohort, but did not replicate ^51^.

Thus, there is increasing evidence of the role of COX in AVS, but it is confounded by the contradictory findings and inconsistent associations. Further research is needed to elucidate the specific roles of AA-COX products in AVS pathogenesis and to explore the shortcomings of COX inhibition.

Future study of dietary modulation to slow AVS progression is also warranted. In many tissues, it has been found that oxylipins can be modified by dietary PUFA ^31,53–55^. It is not yet known whether this is possible specifically in the AV. Our group has, however, previously shown high dietary n-3 PUFA (particularly DHA) to lower prostanoids in the rodent heart ^21^. Whether this can be replicated in the human AV could have potential for future AVS dietary intervention.

A recent study by Artiach et al. (2020) quantified three oxylipins in the AV leaflets of AVS patients and found RvE1 and RvD3 lowered and LtB4 elevated in calcified compared to non-calcified portions of the tissue ^20^. While we did scan for these molecules, they were not above our limits of quantification, thus we cannot comment on them directly. However, we did quantify molecules which are the direct precursors (18-hydroxy-eicosapentaenoic acid and RvE1) or share common precursors (17-hydroxy-docosahexaenoic acid and RvD3; 5-hydroxy-eicosatetraenoic acid and LtB4) and can be considered activation markers. Our finding that 18-hydroxy-eicosapentaenoic acid, 17-hydroxy-docosahexaenoic acid, and 5-hydroxy-eicosatetraenoic acid each increased with AVS conforms with the previous study’s result for LtB4, but not RvE1 and RvD3. The discrepancy may be due to down-regulation of enzymes downstream of the oxylipins we quantified or due to sampling; they compared calcified to non-calcified portions from the same valve, while we homogenized whole leaflets.

In addition to analyzing the oxylipin profile in AV tissue, we also measured plasma oxylipins. We found that the plasma oxylipin profiles were distinct between a healthy control population and patients with AVS. This suggests that the differences in oxylipin profiles between the control and severe AVS groups were more reflective of systemic differences in the two populations than in the AV tissue. The lack of correlation between plasma oxylipins and AVS severity does suggest that changes in oxylipin profiles in AV tissue are not simply a result of systemic inflammation but are specific to the pathophysiology of AVS. This underscores the importance of studying oxylipins directly in AV tissue to understand the underlying mechanisms of AVS. We also acknowledge the limitations in the plasma oxylipin analysis. The control population was considerably younger and had lower rates of smoking and medication use. Also, there was a lack of mild AVS patients (n=5), so it is possible that a modest group separation could have occurred with a greater number of patients.

Sexual dimorphism of oxylipin levels has been previously observed in various animal tissues and human plasma ^21,31,56^. There are also established sex-related differences in AVS outcomes ^57,58^. Interestingly, in this study there were no statistically significant sex-related differences in valvular oxylipins or plasma, and sex was therefore dropped as a factor from the ANOVA models. Similarly, the previous studies examining AV leaflet oxylipins and their synthesizing enzymes did not report sex effects despite sampling males and females, although it is unclear if they considered testing for sex-effects ^16–20^. The lack of sex-effects in this and previous studies may be due to the age of the study participants, in part due to changing sex hormones in older adults ^59,60^.

The present study provides evidence for the presence of elevated prostanoid levels with AVS severity. These findings suggest that prostanoids play a role in the pathophysiology of AVS and may represent a potential target for novel interventions. Further studies are needed to confirm the causative relationship between elevated prostanoids and AVS, and to explore the specific mechanisms underlying this association. Additionally, future research could also evaluate the potential therapeutic benefit of targeting prostanoid signalling in individuals with AVS. Our findings provide novel insights into the role of oxylipins in the pathophysiology of AVS and suggest that targeting oxylipin pathways may be a promising avenue for developing new therapies for this debilitating cardiac condition.

## Non-standard Abbreviations and Acronyms

AA: arachidonic acid
AV: aortic valve
AVS: aortic valve stenosis
COX: cyclooxygenase
DHA: docosahexaenoic acid
EPA: eicosapentaenoic acid
LOX: lipoxygenase
Lt: leukotriene
MPG: mean transvalvular pressure gradient
PCA: principal component analysis
PG: prostaglandin
PUFA: polyunsaturated fatty acids
Rv: resolvin
TX: thromboxane
VIC: valvular interstitial cell

## Acknowledgements

We would like to thank Tanja Winter for support with the oxylipin analysis and Negar Atefi for performing the protein assay. Also, we are grateful to all the patients whose participation made this study possible.

## Supplemental Material

Supplemental Tables 1-6

Supplemental Figures 1-2

